# Individual variation in the dear enemy phenomenon via territorial vocalizations in red squirrels

**DOI:** 10.1101/445767

**Authors:** Jack G. Robertson, Stan Boutin, Murray M. Humphries, Ben Dantzer, Jeffrey E. Lane, Andrew G. McAdam

## Abstract

Territoriality arises when the benefits of resources exceed the costs of defending them. The dear enemy phenomenon, where familiar territorial neighbours refrain from intruding on one another and mutually reduce their defensive efforts, allows for reduction of these costs but requires discrimination between conspecifics. We hypothesized that territorial vocalizations in red squirrels (*Tamiasciurus hudsonicus*) are used for this discrimination. We performed a speaker replacement experiment where red squirrels (n = 41) were temporarily removed from their territories and replaced with a speaker broadcasting their own call, an unfamiliar call, or silence. Contrary to our prediction, there were no differences in overall intrusion risk among our three playbacks, but the identity of intruders did vary. Existing variation in familiarity within territorial neighbourhoods should be considered, rather than the binary classification of familiar or stranger, when studying dear enemy effects. We also discuss the variable importance of silence in acoustic territorial populations.

## Introduction

In populations with intraspecific competition for resources, territoriality as a social structure can arise when individuals benefit from exclusive access to a resource. Resource defense can constitute a substantial portion of a territory owner’s time and energy budget (Puckett & Dill, 1985; Heinemann, 1992), but territoriality is only adaptive when the fitness benefit from the defended resource exceeds the costs of defending said resource from conspecifics (Carpenter & MacMillen, 1976). Furthermore, the net benefit to territory owners can be maximized by reducing defensive costs where possible (Rosell, Gundersen, & Le Galliard, 2008).

Plasticity in territorial behaviour allows for reduction of unnecessary defensive costs. Territory owners across a broad range of species discriminate between conspecifics based on familiarity, known as the dear enemy phenomenon (Fisher, 1954). The dear enemy phenomenon refers to the tendency of territory owners to respond more antagonistically to unfamiliar individuals than familiar territorial neighbours. There are two main predictions of the dear enemy phenomenon: familiar neighbours refrain from intruding on one another, and also reduce their territory defense (Temeles, 1994). This phenomenon likely arises between neighbours as a form of conditional reciprocity, whereby territory owners form ‘agreements’ to tolerate one another and both benefit through a decreased defensive effort (Axelrod & Hamilton, 1981). Individuals that break this relationship and intrude on familiar neighbours incur a cost through increased intrusions on their own territory (Akçay et al., 2009). The dear enemy phenomenon has been demonstrated across taxa, in birds (eg. Wei et al. 2011), mammals (Rosell & Bjørkøyli, 2002), amphibians (Jaeger, 1981), fish (Leiser & Itzkowitz, 1999), reptiles (Qualls & Jaeger, 1991), and crustaceans (Booksmythe, Jennions, & Backwell, 2010).

Most studies of the dear enemy phenomenon test the central predictions by exposing a territory owner to a cue from a neighbouring individual or a stranger and comparing the intensity of the owner’s responses. These experiments are often simple to perform and the pairwise comparison can demonstrate discrimination by the territory owner, but are not directly testing the theory in a territorial context. Such studies also depend on the researchers knowing which identity cue is relevant for their species.

### Individual recognition

The dear enemy phenomenon depends on the ability of individuals to recognize the familiarity of conspecifics in some way. For this to occur, there must be a detectable signal of familiarity between individuals. Identity is typically communicated via the dominant sensory modality for that species, so that birds tend to use songs (Briefer et al., 2008) or calls (Hardouin, Tabel, & Bretagnolle, 2006) but not olfactory cues to recognize conspecifics. Most mammals use scent cues, particularly those where vision or hearing are reduced (i.e. subterranean rodents; Zenuto, 2010), but there is evidence of individually unique vocalizations in mammalian species where acoustic communication is important (Koren & Geffen, 2011). Most avian studies of the dear enemy phenomenon use vocalizations as the presumed cue of individual identity (e.g. Briefer, Rybak, & Aubin, 2008), while most studies of territorial mammals use scent cues (e.g. Müller & Manser, 2007). In experimental tests of the dear enemy phenomenon, the choice of cue is based on the researcher’s hypothesis of how individuals of that species recognize one another. Positive results provide evidence that individuals can discriminate one another, but a finding of no difference between familiar and unfamiliar cues does not necessarily mean the population does not demonstrate the dear enemy phenomenon. Rather, territory owners might be using different information than that provided during the experiment to adjust their behaviour. Our understanding of the dear enemy phenomenon could be improved by testing multiple or atypical modes of recognition within species to determine whether previous findings are biologically meaningful rather than artifacts of experimental design.

### Red squirrels

North American red squirrels (*Tamiasciurus hudsonicus*) are solitary rodents that defend exclusive territories. In the northern boreal forest, these territories are centred on hoards of their primary food source, white spruce (*Picea glauca*) cones, known as middens (Boutin & Schweiger, 1988). A territory with cached resources is required for overwinter survival (Larsen & Boutin, 1995). Juveniles generally acquire a territory before their first winter and remain on the same territory throughout their life. Adult dispersal to new territories is rare, but breeding females may move to another territory and bequeath their own midden to one of their juveniles (Price et al., 1986; Berteaux & Boutin, 2000). Red squirrels primarily defend their territories through vocalizations known as rattles, and direct physical encounters are rare (Smith, 1968; Dantzer et al., 2012). The bioacoustic measures of these calls, such as fundamental frequency and pulse rate, are consistent within individuals (Wilson et al., 2015). Rattles thus provide social information that could be used for individual recognition and discrimination, as red squirrel rattles are individually distinctive (Digweed, Rendall, & Imbeau, 2012; Wilson et al., 2015).

Red squirrel territorial defense is plastic in response to local density (Dantzer et al. 2012), and to the identity of the individuals in their local neighbourhood (Siracusa et al. in review). The dear enemy phenomenon is well demonstrated for this species: squirrels that have been territorial neighbours for longer periods of time are less likely to intrude on each others’ territories (Siracusa, Boutin, et al., 2017), and red squirrels increase their territorial defence behaviour toward unfamiliar individuals (Siracusa et al., in review). Increased territorial defense can be costly in this species in both time and energy (Stuart-Smith & Boutin, 1994), so squirrels in relatively familiar neighbourhoods should not expend maximal effort in defending their territories.

The territorial function of rattles has been empirically demonstrated with a speaker replacement experiment where broadcasting the owner’s rattle reduced the risk of intrusion compared to silence (Siracusa, Morandini, et al., 2017). Behavioural observations of squirrels found that the scale at which the social environment best predicts squirrel behaviour is 150 m around the focal territory (Dantzer et al. 2012), similar to the reported maximum audible distance of a rattle, 130 m (Smith 1978). Red squirrels are sensitive to the acoustic environment around their territory, but rattles may convey more information than just the presence or number of neighbouring conspecifics.

Although the dear enemy phenomenon has been clearly shown for this species, the cue used to recognize the familiarity of conspecifics and adjust behaviour accordingly is not known. Tests of the dear enemy phenomenon in mammals commonly use scent cues (Rosell & Bjørkøyli, 2002; Raynaud & Dobson, 2011; Monclús, Saavedra, & de Miguel, 2014), and red squirrels are able to discriminate conspecifics by scent in captivity (Vaché, Ferron, & Gouat, 2001). This mode of discrimination would be useful for nearest neighbours, but would require squirrels to venture off territory to obtain social information from distant neighbours. Acoustic signals would allow for red squirrel territorial interactions to occur over long distances while individuals remained on their own territories. Given the importance of rattles in territoriality (Dantzer et al. 2012) and the individual specificity of these calls (Wilson et al. 2015), an acoustic mode of discriminating familiarity in red squirrels seems probable. Directed playback studies have found that red squirrels are more likely to respond to playback of unfamiliar rattles than those of neighbours (Price, Boutin, & Ydenberg, 1990), and less likely to respond to rattles of close kin than of unrelated individuals (Shonfield et al. 2017), but these results have been variable (Wilson et al. 2015) and statistical power is limited by the binomial response. We used a speaker replacement experiment in an attempt to investigate the variation in these findings.

### Hypotheses and predictions

Our objective was to investigate whether red squirrels recognize familiarity via vocalizations, facilitating the dear enemy phenomenon in this species. We hypothesized that discrimination between the vocalizations of familiar and unfamiliar conspecifics underlies the dear enemy phenomenon in red squirrels. We tested this hypothesis using a speaker replacement experiment, temporarily removing territory owners and replacing them with a speaker broadcasting their own rattle, the rattle of an unfamiliar individual, or silence. If vocalizations are the cue used to discriminate between conspecifics, then broadcasting an unfamiliar rattle from a given territory should be perceived as a new, unfamiliar owner by neighbouring squirrels.

First, we predicted that there would be a higher risk of intrusion by neighbours when broadcasting an unfamiliar rattle compared to the owner’s rattle; because squirrels refrain from intruding on familiar individuals, this unfamiliar rattle should be less effective at deterring neighbours from intruding than the territory owner’s rattle. Silence should also lead to a higher risk of intrusion by neighbours than the owner’s rattle, as shown by Siracusa, Morandini, et al. (2017).

Second, as per the dear enemy phenomenon, simulating an unfamiliar territory owner should lead to higher defensive effort by neighbouring squirrels. We predicted that rattling rates in the surrounding acoustic neighbourhood would be higher when broadcasting an unfamiliar rattle than when broadcasting the owner’s rattle during our temporary removal experiments.

## Methods

### Study system

This study was part of the Kluane Red Squirrel Project, a long term study of a wild population of red squirrels within Champagne and Aishihik First Nations Traditional Territory along the Alaska Highway in the southwest Yukon (61° N, 138° W), which has been running since 1987 (McAdam et al., 2007). The study site is boreal forest dominated by white spruce, the cones of which are cached in middens near the centre of each squirrel’s territory.

As part of this project, every red squirrel living within several ~40ha study grids was tagged with unique alphanumeric metal ear tags and monitored throughout its life. Coloured wires and pipe cleaners were threaded through the metal ear tags to allow identification of individuals from a distance. Territory ownership was determined through biannual censuses of the population every spring and fall based on live trapping and behavioural observations, so that the location and duration of ownership was known for every individual in the population.

We selected a semi-random sample of adult male squirrels (n = 42) from two control study grids (KL and SU) as our focal individuals, each territory separated by > 60m, to reduce confounding effects of neighbours between trials. We did not use female squirrels both due the ethical concerns of removing a mother from her pups during lactation, as well as the potential for maternal protection to affect the intensity of territory defense independent of the social information of interest.

### Rattle collection and processing

We recorded rattles from each of our focal individuals to use in the speaker replacement trials. Our speaker replacement experiment had a repeated measures design, where each of the 42 individuals were temporarily removed from their territory three times, and replaced with a speaker broadcasting their own rattle, an unfamiliar rattle, or silence in a randomized order. Each individual’s recorded rattles were used twice, once at their own territory as the owner treatment and once at the territory of an individual on the other study grid as the unfamiliar treatment.

We deployed an audio recorder (Zoom Corporation^®^ H2N audio recorder, Tokyo, Japan) on the midden of each of our focal squirrels and recorded for 24 hours. We then went through these recordings and selected the three highest quality recordings of rattles from each individual. Owner rattles were distinguished from neighbours by amplitude, as neighbouring squirrels would not be rattling on the owner’s midden. This approach has been previously shown to reliably identify the calls of owners (Siracusa et al. in review). These three rattles were extracted and normalized to the same amplitude for every individual using Avisoft-SAS Pro software (Avisoft Biacoustics), but were otherwise left unmanipulated. We combined these three rattles in a 21- minute audio file with seven minutes of silence separating each rattle; one rattle every seven minutes is the average natural rate for this population (Dantzer et al., 2012).

### Speaker replacement experiment

We trapped focal individuals using Tomahawk traps (Tomahawk Live Trap, Tomahawk, WI, USA) and temporarily removed them from their midden. Squirrels were placed in a modified box (41 × 17.5 × 19 cm) and provided with a small amount of peanut butter and a slice of apple for food and hydration (Donald & Boutin, 2011). The box was placed in a sheltered location away from other squirrel middens, and disinfected with isopropyl alcohol between removals. We conducted all trials during May – August 2017 between 7 am – 12 pm, the main activity period of squirrels during the summer months (Studd et al., 2016), and avoided days with precipitation or high winds, as squirrels tend to be less active during these weather conditions (Williams et al., 2014).

Once the owner was removed from its territory, we set up an SME-AFS field speaker (playback range 0.1 – 22.5 kHz, Saul Mineroff Electronics) on the centre of the midden. The speaker played one of the two audio treatments – the owner’s rattle or an unfamiliar rattle – or was left inactivated for silent trials. We used a silent control as opposed to a noise control because other sounds in the environment such as bird calls could communicate additional information (Randler, 2006), and we were interested in how neighbouring squirrels would respond to the absence of any acoustic information coming from the focal territory. Rattles were broadcast from this speaker at natural levels of 65–75 dB (Shonfield et al., 2017) measured 2 m from the speaker using a digital sound level metre (RadioShack model 33-2055A). We deployed a Zoom recorder on the focal midden, to record the rattling behaviour of neighbouring individuals throughout the removal. We also manually noted the time and approximate location of any audible neighbour rattles during each trial, as the Zoom recorder could not identify the direction from which each rattle was emitted.

Each squirrel was temporarily removed three times, with the treatments performed in randomized order for each individual, separated by 21 - 48 days (median = 28). As dispersal between the two study grids is low, and no squirrels were within 130 m of a squirrel on the other study grid, we paired squirrels so that each playback file was used twice, once at their own territory and once as the unfamiliar treatment at a territory on the other study area. The playback audio files were given 5-digit numerical names so that the identity of the rattle was not known when playing the files and collecting data in the field. The audio treatments during temporary removals were thus partially blind, because the researcher performing the removals did not know if it was the owner or an unfamiliar rattle playing, but the silent trials were identifiable.

We observed the focal territory from >10 m away and recorded the time and identity of the first squirrel to step over the edge of the midden, or travel through trees to cross this same threshold. The edge of the midden was defined by the extent of visible cone bracts on the ground. Removals ended after the first intrusion, or if no intrusion occurred, after a maximum latency of two hours. Upon completion of the trial, the owner was returned to the midden and released, and the speaker was removed. This research was approved by the University of Guelph Animal Care Committee (AUP 1807).

When performing a temporary removal experiment, the owner is prevented from calling to defend its territory; the treatment is effectively removing their territorial vocalizations from the acoustic environment, and replacing these vocalizations at a hypothesized rate with a speaker. To quantify the difference between our silence and playback treatments, it is important to know actual vocalization rates in the study population during the experiment. To characterize the social context of our population in which we conducted our study, we deployed Zoom recorders on the middens of other squirrels on our study grids (n = 42) as described above in *Rattle collection and processing*. These audio recordings were analyzed as described in Siracusa et al (in review) (modified to include the hours of 08:00 – 13:00 rather than 07:00 – 13:00 due to reduced temporal coverage in 2017) to obtain an average number of rattles per individual throughout the morning active period during which we conducted our speaker replacements.

### Long term data

As part of the Kluane Red Squirrel Project, the age, sex, and history of territory ownership for every tagged individual in the population was known. We identified every squirrel living within acoustic range of the focal individuals during the experiment, based on the 130 m range of red squirrel rattles (Smith, 1978). By identifying the earliest census in which the focal squirrel and a neighbour occupied their respective territories, we calculated pairwise familiarity with each neighbour as the number of days that those two squirrels occupied neighbouring territories. Neighbourhood density, age and sex of neighbours, and the distance of each neighbour from the focal midden were also obtained from our existing long-term data.

### Statistical analysis

We tested the effect of the rattle playback on the risk of intrusion using a Cox proportional hazard mixed effects model. This model works well with censored data, where the time to an event is recorded but in some trials the event never occurs. The binary response (did an intrusion occur?), and the latency to the event (how long did it take for the intruder to appear?), which has a maximum value of 120 min, were incorporated together as a single response known as a hazard function. We included audio treatment as a three-level categorical predictor, and because there were repeated measures for the same individual, we also included a random effect of owner ID to account for variation in intrusion risk among neighbourhoods.

We analyzed neighbourhood rattling rate (number of rattles/min from all neighbours) during each removal using a linear mixed effects model, testing for an effect of audio treatment while also incorporating day of year, local neighbourhood density and familiarity, and a random effect of owner ID.

As neighbourhood-level analyses found no evidence of rattle identity on intrusion risk (see *Overall intrusion risk* results below), we performed two sets of post hoc analyses. First, we added several other neighbourhood covariates that are known to affect intrusion probability and intensity of defense (Siracusa, Boutin, et al., 2017) to the models of hazard of intrusion and rattling rate. We included local neighbourhood density, average neighbourhood familiarity, study grid, and day of year in each model, and we interacted audio treatment with each of these in turn to test whether responses to different playback types were dependent on one of these other factors.

Next, based on Siracusa, Boutin, et al. (2017) which found that unfamiliar neighbours were more likely to intrude than familiar neighbours within the same neighbourhood, we modelled the individual hazard of intrusion of each neighbour intruding at each temporary removal. For each speaker replacement trial, all squirrels living within 130 m of the removal territory were considered potential intruders and coded as intruding (1) or not intruding (0) for each temporary removal. We used a Cox proportional hazard model as for the neighbourhood-level model above, but now testing which of the neighbours intruded, rather than whether any intrusion occurred at the focal territory. As we ended removals after the first intruder appeared, when one neighbour intruded it prevented us from detecting intrusions by other neighbours. The Cox proportional hazard model tests whether an event occurs within a given maximum possible timespan; when an intrusion occurred, this maximum latency was less than 120 minutes for the other neighbours. Thus, if no squirrel intruded, we scored all neighbours as 0 and 120 min was the maximum latency; if a neighbour intruded after 45 min, we scored this neighbour as a 1 while all other neighbours were 0, and all squirrels had the same maximum latency of 45 min. For trials where a non-neighbouring squirrel intruded, the maximum latency for all neighbours was the time to this intrusion, but we scored all individuals as 0 (not intruding) because the intruder was not a member of this neighbourhood.

To investigate which characteristics might predict how neighbours responded to the audio treatments, we included day of year, neighbour sex, and within-neighbourhood standardized measures of neighbour age, pairwise familiarity with the removed owner, and distance from focal midden as fixed effects, and a random effect of trial ID. We tested for significant interactions with playback type for each of these in turn.

All statistical analyses were conducted using R version 3.3.3 (R Core Team 2017), using the packages coxme (version 2.2-5, Therneau, 2015), lmerTest (version 2.0-33, Kuznetsova, Brockhoff, & Christensen, 2016), and visreg (version 2.4-1, Breheny & Burchett, 2017). Reported estimates are means ± SE.

## Results

We performed a total of 126 temporary removals among 42 male squirrels. Some trials had to be discarded due to weather (rain during the trial) or speaker malfunctions; excluding these, there were 115 removals across the three playback types among 41 male squirrels. Of these 115 trials, there were four (one silent, one owner playback, two unfamiliar playback) in which an intruder appeared within three minutes of the removal starting. In all stimulus tracks, the first rattle in the audio track played after three minutes of silence, so in these removals an intrusion occurred prior to the first possible rattle in the recording. These trials cannot be considered as either playback, and rather than reclassifying them as silent trials – which would substantially increase the overall risk of intrusion for the silent treatment – they were removed from the analysis. Thus, the models described below include 111 temporary removals among 41 individuals: 35 owner playback, 34 unfamiliar playback, and 42 silent trials.

There were 19 trials of these 111 in which an intrusion occurred, but the intruding individual was identified as a squirrel living outside the 130 m radius around the removal territory. These squirrels cannot be considered to have experienced the speaker treatment equivalently to neighbouring squirrels to the removal territory. The dear enemy phenomenon would only predict differing risks of intrusion when playing an owner’s call compared to an unfamiliar call if all individuals hearing these rattles broadcast were familiar with the territory owner. For squirrels outside the social acoustic neighbourhood, there is no reason to predict discrimination between these rattles because both calls are unfamiliar. Additionally, it is unclear to what extent these squirrels would have heard any rattles being broadcast as their territories lie outside the acoustic range of the speaker. Thus, all results presented below include these 19 trials redefined as not having an intrusion but with a maximum latency of when the non-neighbour intrusion occurred.

Local density for our focal squirrels was 2.65 ± 0.12 squirrels/ha and familiarity was 495 ± 31 days. Local densities on KL (3.11 ± 0.15 squirrels/ha) were higher than those on SU (2.22 ± 0.11; *t* = 4.72, df = 39, *p* < 0.0001), but neighbourhood familiarity did not differ (*t* = 0.54, df = 39, *p* = 0.59). Squirrels on KL rattled more than those on SU during a five-hour period in the morning (32.1 ± 9.2 vs. 13.5 ± 2.1 rattles; *t* = 2.25, df = 40, *p* = 0.03). The overall average was 21.5 rattles over five hours, corresponding to one rattle every 14 minutes.

### Overall intrusion risk

There was no overall effect of the playback type on the risk of intrusion by neighbours (Figure 1). The overall probability of intrusion was 24 %, and the proportion of trials with an intrusion did not vary between treatments (Figure 2), nor did the average latency to an intruder (owner = 43.7 ± 7.7 min, silence = 46.1 ± 7.7 min, unfamiliar = 59.5 ± 7.9 min; overall = 49.1 ± 4.6 min). Including all intrusions, or only those from within the social neighbourhood, did not change the interpretation of the playback effect.

**Figure 1.**
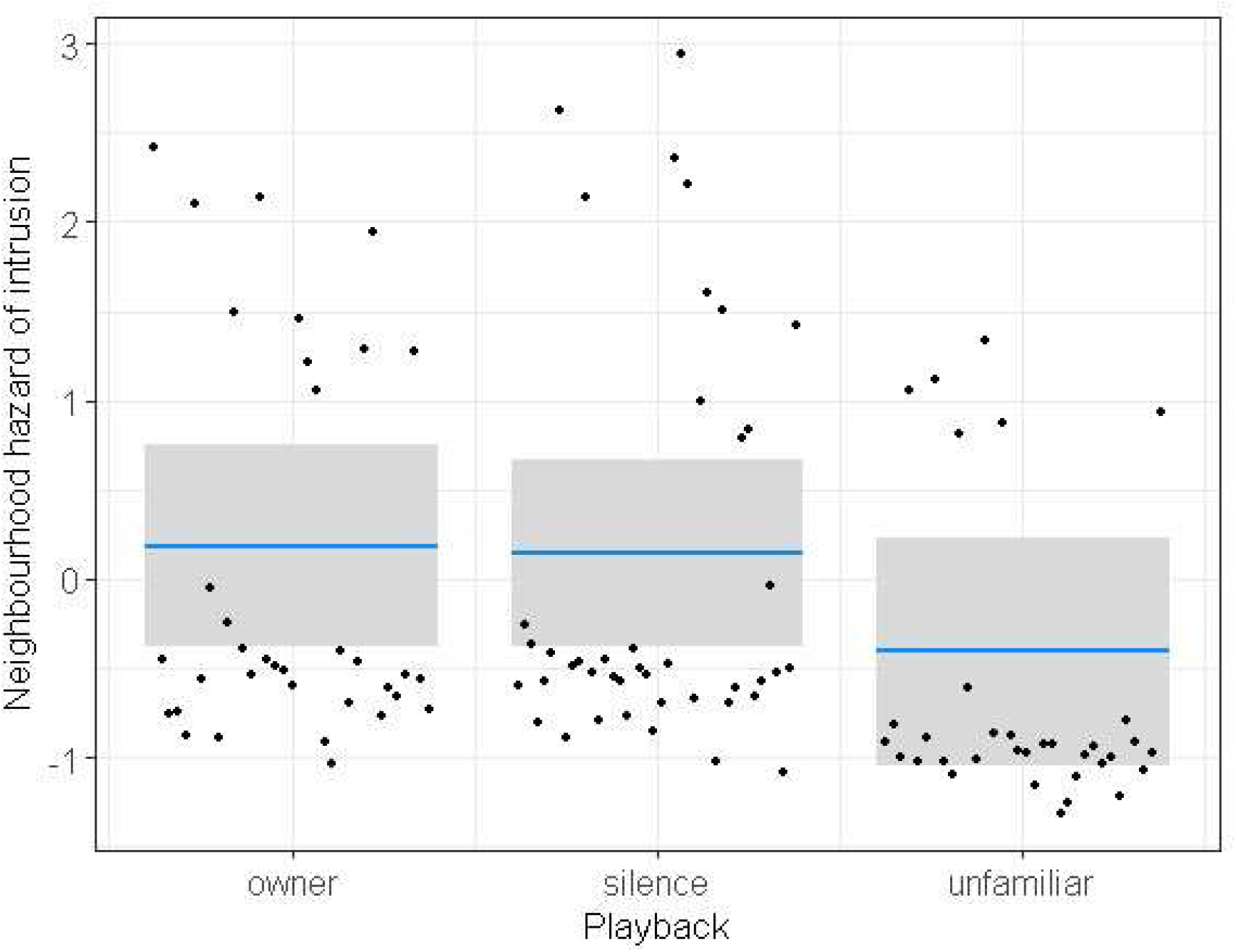
Cox proportional hazard model of intrusion by neighbours, incorporating whether or not an intrusion occurred with the latency to that intrusion event. There were no differences in intrusion hazard between owner playback and silence (z = −0.15, *p* = 0.88) or owner and unfamiliar playbacks (z = −1.48, *p* = 0.14).

**Figure 2.**
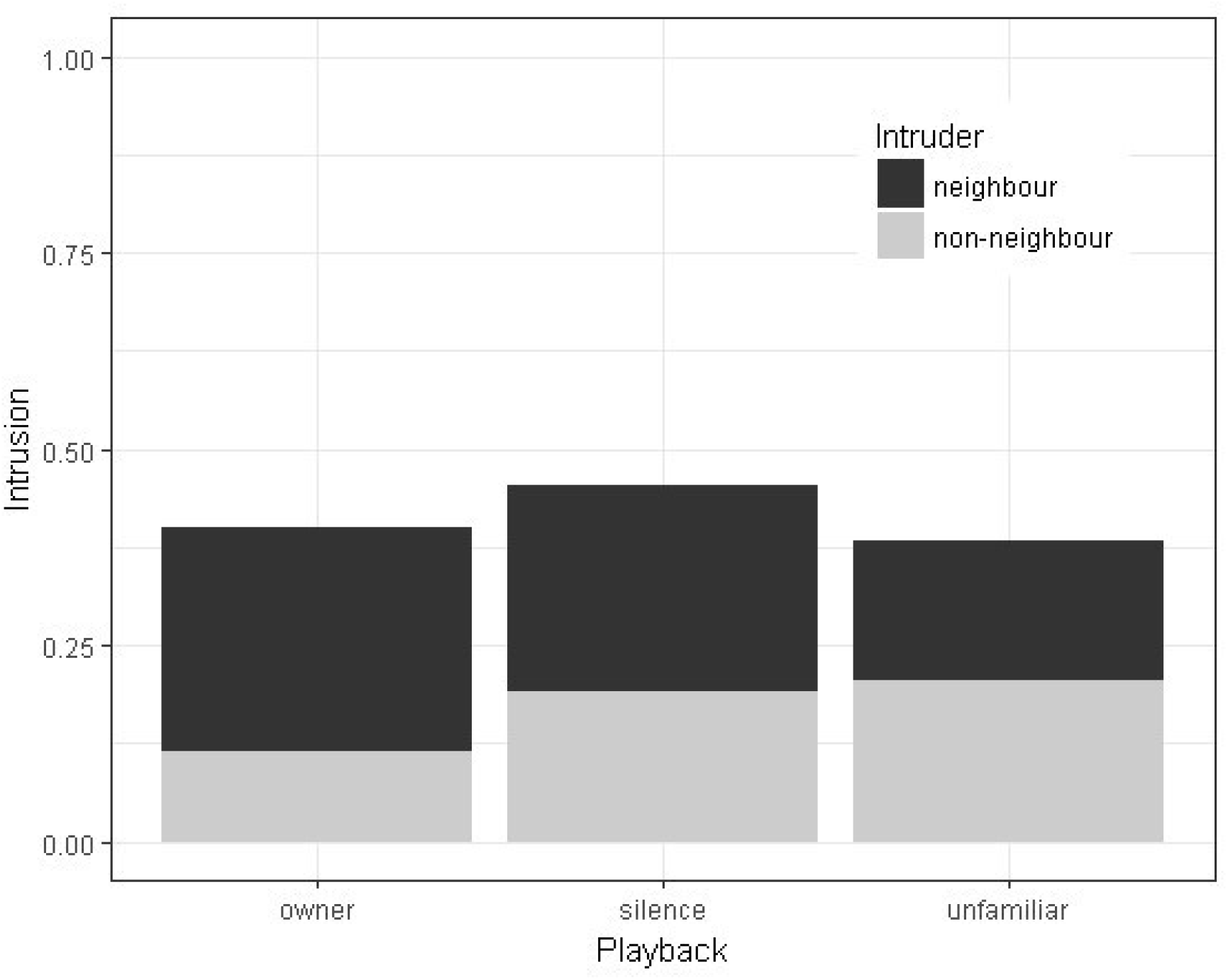
Proportion of trials across three playback treatments where a neighbour (black) or non neighbouring (grey) squirrel intruded during a 2-hour temporary removal. There were no significant differences between playbacks when considering all trials, only neighbour intrusions, or only non-neighbour intrusions.

It is unlikely that censoring trials upon the first intrusion affected these relationships. If either non-neighbour intrusions or those from within the social neighbourhood happened earlier, it is possible that detecting intruders from a different source was prevented by ending the trial upon seeing the first squirrel. However, in about 60% of all trials, there were no intruders at all; if there was some sort of masking effect preventing detection of both types of intruders at a given territory, there should be fewer removals with no intruder at all. Additionally, there was no difference between the average latencies of intrusions coming from within the neighbourhood (45.9 ± 5.8 min) and those from non-neighbours (53.8 ± 7.4 min) (*t* = 0.85, df = 44, *p* = 0.40), so it is unlikely that censoring trials when the first intruder appeared prevented detection of a later intrusion.

### Neighbourhood rattling rates

The number of rattles/min heard by observers during the removals was highly right skewed (mean = 0.54, median = 0.46, range = 0 – 2.49 rattles/min). This response was, therefore, log10(x + 0.1) adjusted prior to analysis (+ 0.1 as three trials had zero rattles heard). Contrary to our second prediction, rattling rate in the neighbourhood did not vary among the three treatments (*F*_2,76_ = 0.267, *p* = 0.77). Although trials varied in length when intrusions occurred, the duration of the trial did not affect the observed rattling rate (*t* = −0.27, df = 109, *p* = 0.79).

### Post hoc analyses

In the Cox proportional hazard model of neighbourhood intrusion risk, none of the other neighbourhood factors had a significant effect on the risk of intrusion (familiarity: z = 1.15, *p* = 0.25; date: z = 0.55, *p* =0.66; grid: z = 1.16, *p* =0.25; density: z = 0.54, *p* = 0.59). Interacting playback type with each of these covariates in turn, also provided no evidence that the effect of the speaker on intrusion hazard was dependent on any of these factors.

Although these neighbourhood characteristics did not predict the risk of intrusion, some did have an effect on rattling rates during the temporary removals. Unsurprisingly, local density had a positive effect on neighbourhood rattling rate; with more neighbours around the removal midden, there were more audible neighbour rattles (*t* = 2.46, df = 109, *p* = 0.01). When accounting for the effect of local density, there were also more rattles on KL grid than on SU (*t* = 2.93, df = 109, *p* = 0.004), suggesting that there are underlying differences in the behaviour of the squirrels in these areas beyond the differences in population density. Neighbourhood rattling rate also increased with day of year, with more rattles later in the summer (*t* = 2.71, df = 109, *p* = 0.007). Average familiarity around the removal territory did not affect neighbourhood rattling rates (*t* = 1.03, df = 109, *p* = 0.31). As squirrels adjust their rattling behaviour in response to their local familiarity and density (Siracusa et al, in review), we calculated familiarity and density for the 130m radius around each individual neighbour squirrel and included these values in our model. The average of the familiarity and density values for every neighbour around each removal also did not affect the neighbourhood rattling rate observed during the speaker replacements (familiarity: *t* = 0.67, df = 109, *p* = 0.50; density: *t* = 0.68, df = 109, *p* = 0.50). There was no significant effect of playback with the inclusion of these covariates in the model (*F*_2,104_ = 0.37, *p* = 0.69). Interactions between the playback type and each of these factors were tested, but no combinations were significant.

### Individual hazard of intrusion

The second stage of post hoc analyses considered which individual squirrels within the acoustic neighbourhood were more likely to intrude during a given trial. Red squirrel neighbourhoods are not homogenous, and neighbours have varying degrees of familiarity with each focal individual. Our original prediction that there would be more intrusions during unfamiliar rattle playbacks presumed that all neighbours were uniformly familiar with the removed squirrel and would all exhibit dear enemy relationships. However, new neighbours or squirrels that have just joined the social neighbourhood would not have as much familiarity with the focal squirrel being removed, and so might not be expected to discriminate as strongly between the two playback types, leading to less difference between the audio treatments.

Overall, closer neighbours had a higher hazard of intrusion (β = - 0.02, z = −3.50, *p* = 0.0005) and younger neighbours tended to intrude more (β= −0.31, z = −1.39, *p* = 0.16). Neighbour sex did not affect intrusion hazard (z = −0.55, *p* = 0.58). Interacting the standardized familiarity of neighbours with the playback type, the discrimination between owner and unfamiliar playbacks varied depending on how familiar the neighbour was with the focal squirrel (Figure 3). Relative familiarity within the neighbourhood had no effect on individual hazard of intrusion during silence (slope = −0.16 ± 0.34, *p* = 0.63) or unfamiliar playback (slope = −0.07 ± 0.49, *p* = 0.88). However, when broadcasting the owner’s call, unfamiliar neighbours were more likely to intrude, while familiar neighbours refrained from intruding (slope = −1.12 ± 0.39, *p* = 0.004).

**Figure 3.**
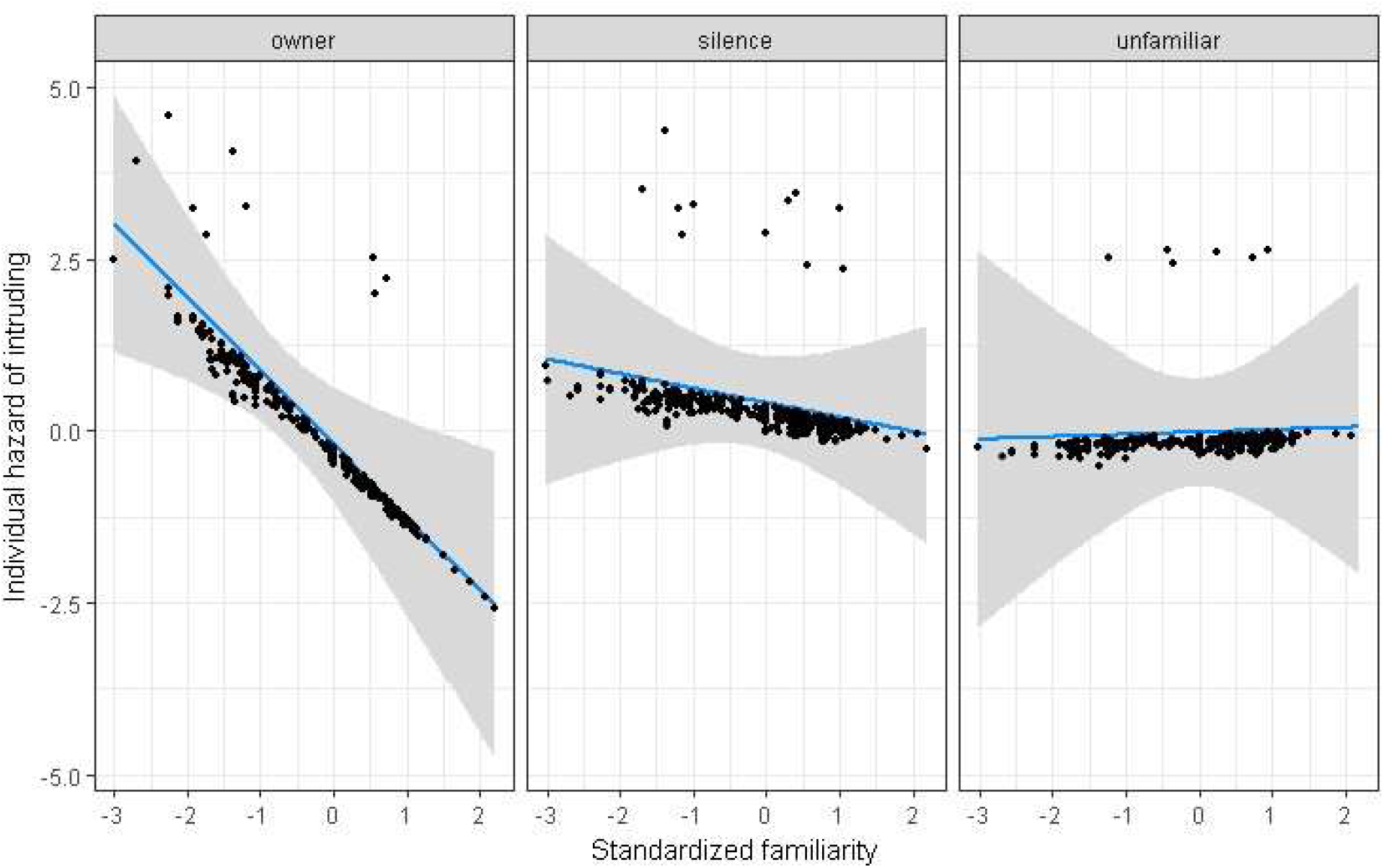
Relative hazard of intrusion by individual neighbouring squirrels during a temporary removal, under three audio treatments. Intrusion hazard is modelled using a mixed effects Cox proportional hazard model, accounting for the latency to an intrusion event and whether or not one occurred within the two-hour temporary removal; a high hazard corresponds to a shorter latency and higher probability of intrusion. There was a higher hazard of intrusion from relatively unfamiliar neighbours when broadcasting the owner’s rattle (β= −1.05, z = −2.82, *p* = 0.005), whereas relative familiarity did not affect the hazard of intrusion during silence (β = -0.16, z = 0.49, *p* = 0.63) or unfamiliar rattle playbacks (β = 0.07, z = 0.15, *p* = 0.88).

The differences in intruders between the three treatments can be elucidated by comparing the relative familiarity of intruders under the three playback types (Figure 4). When the owner’s rattle was broadcast, intruders (*n* = 9, mean familiarity z-score = −1.04 ± 0.44) were less familiar than other neighbours (*t* = −2.85, df = 23, *p* = 0.009), while intruders during silent trials (*n* = 14, z-score = −0.22 ± 0.27) and trials playing an unfamiliar call (*n* = 6, z-score = - 0.02 ± 0.33) were not different from the average familiarity within the neighbourhood (silence *t* = −0.84, df = 23, *p* = 0.41; unfamiliar *t* = −0.03, df = 23, *p* = 0.97). Squirrels intruding when the owner’s rattle was broadcast were marginally less familiar than intruders during the unfamiliar playback (*t* = −1.8, df = 23, *p* = 0.08).

**Figure 4.**
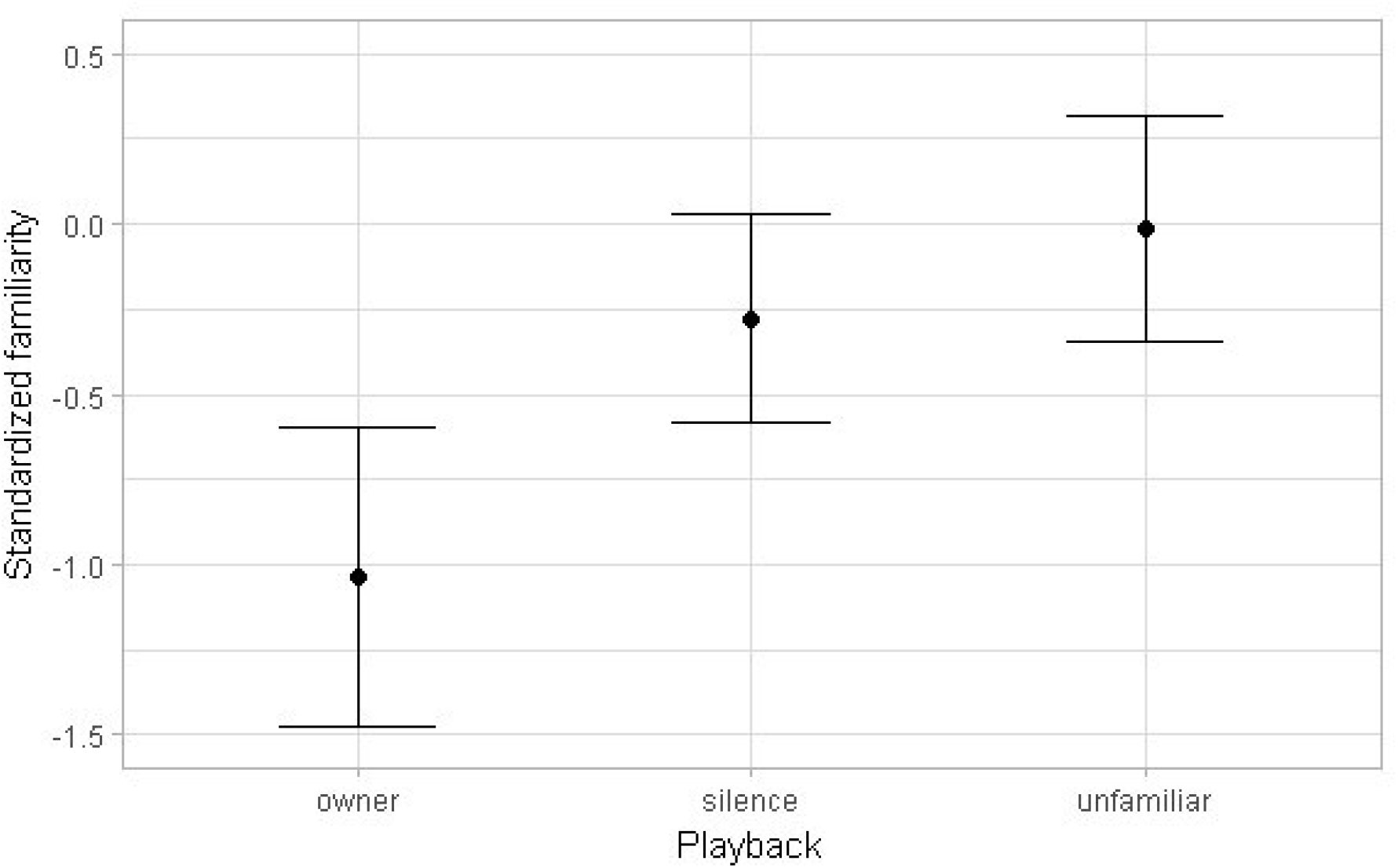
Standardized familiarity (z-scores, relative to neighbourhood) of intruding neighbours during temporary removals when broadcasting the owner’s call (n = 9), silence (n = 14), or an unfamiliar call (n = 6). Standardized familiarity for intruders on the owner’s rattle was different from zero (t = −2.85, df = 23, p = 0.009), but this was not the case for either of the other two groups (silence: t = −0.84, df = 23, p = 0.41; unfamiliar: t = −0.03, df = 23, p = 0.97).

## Discussion

Our a priori prediction that neighbouring red squirrels would intrude less when the territory owner’s rattle was broadcast from its territory as opposed to a broadcast of an unfamiliar rattle was not supported by this study, nor did we find that either rattle reduced the risk of intrusion relative to silence. Similarly, rattling rate from surrounding territories was unaffected by the identity of the rattle played. However, post hoc analyses revealed that neighbouring squirrels had differing responses to the owner and unfamiliar rattles based on pre-existing familiarity with the owner: familiar neighbours refrained from intruding during owner playback but did intrude when broadcasting the stranger rattle, whereas unfamiliar neighbours did not discriminate rattle identity as strongly. Differing responses of familiar and unfamiliar neighbours masked any differences in intrusion risk overall at the neighbourhood level, but the identity of intruders differed between playback types (Figure 4). The difference in familiarity of intruders between owner and unfamiliar speaker replacements supports the ability of red squirrels to recognize familiarity in the rattles of conspecifics, but the behavioural responses did not differ as initially predicted.

Most tests of the dear enemy phenomenon define familiarity as a binary of neighbours and strangers. However, familiarity with territorial neighbours is a continuum, and in longer-lived year-round territorial species familiarity can accrue substantially. Average lifespan of red squirrels in this area that recruit into the population as adults is 3.5 years (McAdam et al., 2007), but the highest pairwise familiarity between two red squirrels in this study was 6.2 years. Other squirrels had been neighbours for less than a week prior to the temporary removal: these situations are evidently not equivalent, but both would be categorized as neighbours in a binary model. The lack of overall difference in intrusion risk between the familiar and unfamiliar rattle playback, but the clear difference in response to these treatments based on pre-existing familiarity, corroborates the necessity of considering familiarity in territorial populations as a continuous measure. The available data for territory occupancy from this long-term project allowed us to identify these individually variable responses that were masked when averaging across neighbourhoods. These more nuanced effects of individual familiarity in response to cues from neighbours might similarly obscure dear enemy relationships in other systems where familiarity has been classified as neighbour or non-neighbour. Further studies of the dear enemy phenomenon would benefit from considering individual variation within territorial populations, and the effects this variation can have on observed overall trends.

At first our results appeared to be contrary to previous studies documenting the territorial defense function of rattles (Siracusa, Morandini, et al., 2017) and the effects of familiarity on intrusion risk (Siracusa, Boutin, et al., 2017), but this is not surprising given the natural inter annual fluctuations in food, density, and predators in this population (Boutin et al., 2006; Dantzer et al., 2013; Studd et al., 2015). Overall, intrusion rates in this study were lower than those observed in a previous speaker replacement study (Siracusa, Morandini, et al., 2017). This previous study was conducted in fall 2015, a year immediately following a white spruce masting event in our study area. In a mast year, trees coordinate their production of cones to create a superabundance of resources, followed by several years of very low or zero cones produced so that cone availability varies by several orders of magnitude (Lamontagne & Boutin, 2007). Squirrel populations track this variation, with densities ranging from 0.4 – 3.0 squirrels/ha (Dantzer et al., 2013). Recruitment of juvenile red squirrels is much higher than usual following a mast year (McAdam & Boutin, 2003), resulting in high densities and low familiarities in the subsequent year. With minimal juvenile recruitment in subsequent low cone crop years, density typically declines and average familiarity increases in the years following a mast event. The average local density and familiarity in this study were significantly lower (*t* = −8.41, df = 94, *p* < 0.0001) and higher (*t* = 8.48, df = 94, *p* < 0.0001) respectively than those in 2015 (Figure 5). Although these inter-annual differences could explain the reduction in overall intrusion risk between years, they should not affect the discrimination between familiar and unfamiliar calls, nor the effectiveness of rattle playbacks.

**Figure 5.**
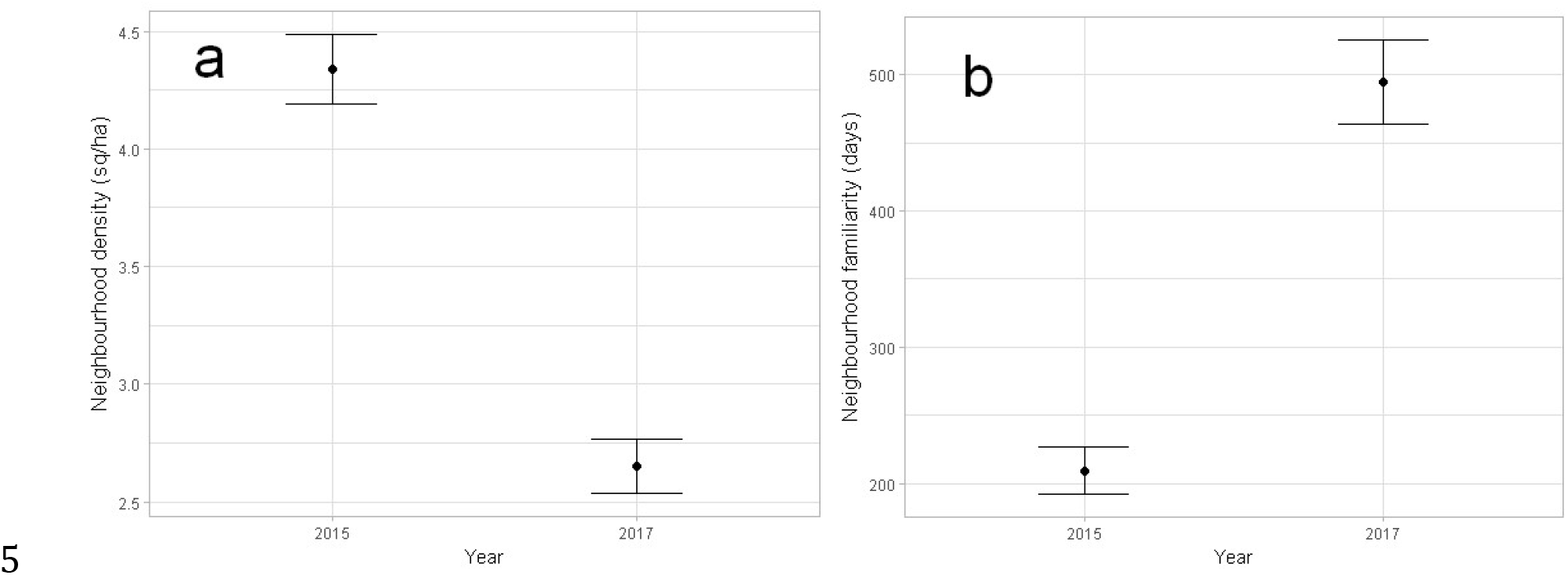
Comparison of (a) local density and (b) familiarity within 130m of focal squirrels used for temporary removal experiments in 2015 (n = 55) and 2017 (n = 41). With two further years of very low cone production since the 2014 mast event, population density has fallen with the lack of new food resources while average familiarity in 2017 has accrued as there was minimal recruitment of juveniles during this period.

The central anomalous finding of this study, in light of the results from Siracusa, Boutin, et al. (2017), is that unfamiliar neighbours failed to intrude during silent trials, but did intrude on the owner’s rattle. Why these unfamiliar squirrels refrained from intruding during silence, but were seemingly attracted by the owner’s rattle, is unclear. Potentially, in a social environment with scarce acoustic information, the presence of the owner rattle attracted these unfamiliar neighbours to intrude, but the lack of information during silence created uncertainty that discouraged these squirrels from intruding. This raises questions of the importance of silence in populations where the primary territorial defense is acoustic.

Siracusa, Morandini, et al. (2017) documented a strong effect of broadcasting the owner’s rattle relative to silence, reducing the risk of intrusion at the same territory by 2.5 times. This is a much stronger effect than that observed in this study, and this discrepancy can likely be explained by year to year variation in both intrusion pressure and ambient rattling rate. Siracusa, Morandini, et al. (2017) conducted their trials in the fall, when neighbourhood rattling rates during removals in this study were highest (*t* = 2.71, df = 109, *p* = 0.007). Under conditions where ambient rattling rate is high, such as low familiarity and high density (as in 2015), the absence of rattles from a territory becomes a meaningful source of information to neighbours that the owner is not actively defending their midden. However, squirrels on the same study grid (KL) rattled 45% less in 2017 than they did in 2015: 46.44 ± 1.8 (n = 85) rattles per morning in 2015 vs. 32.1 ± 9.2 (n = 18) in 2017 (*t* = 2.52, df = 101, *p* = 0.013). Our speaker replacement used the previously reported average rattling rate of 1 call per 7 minutes (Dantzer et al., 2012), whereas the average time between rattles across both grids in 2017 was 14 minutes. A silent 2- hour removal thus corresponds to only 8 or 9 missing rattles, rather than 17 rattles based on the playback rate of one call every seven minutes. This drastically reduces the value of silence, relative to either owner or unfamiliar rattle playback, in determining whether a territory is being defended.

Speaker replacement or targeted playback experiments are often used to test the dear enemy phenomenon or the value of territorial vocalizations more generally. An important component of these studies is the temporal rate at which calls are broadcast from the speaker; this is generally based on reported average vocalization rates for the study species, but as demonstrated in this study, these rates are not fixed values and the speaker may not be simulating typical defensive behaviour in that population. Comparisons of defensive calls relative to silence on a territory are implicitly testing the value of silence as information, which is dependent on the expected amount of acoustic information present in the environment.

Our speaker replacement experiment demonstrated that red squirrels are capable of discriminating familiarity in vocalizations, but did not respond behaviourally as we had initially predicted. Our results differed from previous temporary removal and speaker replacement studies in the population, likely due to interannual differences in social composition and defensive effort. The extensive within-neighbourhood variation in familiarity documented in this system, and the individually variable responses of neighbours to familiar and unfamiliar vocalizations, reinforces the need to consider territorial familiarity as a continuously variable trait within populations, rather than as a binary of neighbours and strangers. We also highlight the importance of silence as information in acoustically territorial populations, and how the relative value of silence depends on expected rates of vocalization.

## Acknowledgements

We thank the Champagne and Aishihik First Nations for providing access to the land on which the study sites for this project were located, in particular Agnes MacDonald and her family for long-term access to her trapline. Analyses and writing for this manuscript were conducted at the University of Guelph, which resides on the ancestral lands of the Attawandaron people and the treaty lands and territory of the Mississaugas of the Credit. E. Siracusa provided invaluable guidance in design and analysis of this study, and D. Wilson was integral to the analyses of audio recordings. Particular thanks to M. Thorpe for her integral assistance in field data collection. Research funding for this project was provided by a Grant in Aid of research from the American Society of Mammalogists (J. Robertson), the Northern Scientific Training Program (J. Robertson), an Ontario Graduate Scholarship (J. Robertson), University of Michigan (B. Dantzer), and the National Sciences and Engineering Council of Canada (J. Robertson, A.G. McAdam, S. Boutin, M.H. Humphries, J.E. Lane). This is publication 91 of the Kluane Red Squirrel Project.

